# SPATEs promote the survival of *Shigella* to the plasma complement system upon local hemorrhage and bacteremia

**DOI:** 10.1101/2023.11.08.565994

**Authors:** Lorine Debande, Ahmad Sabbah, Laurianne Kuhn, Patryk Ngondo, Antonin C André, Béatrice Roche, Matthieu Laborde, Victoria Cantalapiedra-Mateo, Tamou Thahouly, Ana Milinski, Laurent Bianchetti, Christine Allmang-Cura, Magali Frugier, Benoit S Marteyn

## Abstract

*Shigella* spp. are the causative agents of shigellosis, which remains a leading cause of death in children under the age of five. Shigellosis is characterized by fever and results in hemorrhagic diarrhea; in more severe cases, *Shigella* bacteremia has been reported. These clinical features strongly suggest that *Shigella* survive exposure to plasma, although this has not yet been studied at the molecular level. In this report, we confirmed in a guinea pig model of shigellosis that local hemorrhages were induced by *S. flexneri* 5a and *S. sonnei*, and we demonstrated that *Shigella* reached mucosal CD31+/CD34+ blood vessels during the late stages of infection and further disseminated in the bloodstream. These results confirmed the exposure of *Shigella* to plasma components within the hemorrhagic colonic mucosa and in the bloodstream. We demonstrated that all the tested *Shigella* strains survived plasma exposure *in vitro*, and we showed that Serine Protease Autotransporters of Enterobacteriaceae (SPATEs) are essential for *Shigella* dissemination within the colonic mucosa. We have confirmed that SPATEs are expressed and secreted in poorly oxygenated environments encountered by *Shigella* from hypoxic foci of infection to the bloodstream. We further demonstrated that SPATEs promoted *Shigella* survival in plasma, by cleaving complement component 3 (C3), thereby impairing complement system activation. We have shown here that the ability of *Shigella* to survive plasma exposure is a key factor in its virulence, both within primary foci and systemically.

**Significance Statement:** In this study we aimed to better understand the significance of the ability of *Shigella* to survive plasma exposure, as we observed that non-pathogenic *E. coli* rapidly lysed upon exposure. Indeed, we reported that *Shigella* was already exposed to plasma components within the colonic mucosa, as we reported in a guinea pig model of shigellosis that hemorrhages were induced, that were associated with local diffusion of plasma components in the infected colonic mucosa. *Shigella* was obviously exposed to plasma during bacteremia. The ability of *Shigella* to survive in plasma has not been previously reported. Here we have shown, first, that *Shigella* was able to divide and grow in the presence of human plasma, and second, we found that SPATEs played a central role in this process by impairing with the activation of the complement system.

## Introduction

Shigellosis, or bacillary dysentery, is caused by *Shigella* spp., a genus of pathogenic enterobacteria that specifically infect and colonize the human colon. Four species of *Shigella* have been identified: *S. flexneri, S. sonnei, S. dysenteriae*, and *S. boydii*. Millions of cases of shigellosis are still reported worldwide each year, resulting in hundreds of thousands of deaths in developing countries (1-4). Symptoms of shigellosis include severe hemorrhagic diarrhea, vomiting, fever, abdominal pain and dehydration. Evidence of bloody and mucoid diarrhea in shigellosis patients is caused by local hemorrhage within the colonic mucosa, associated with endothelial damage. This clinical aspect of shigellosis has long been recognized, but the impact of the presence of plasma components with strong antimicrobial activity within the colonic mucosa on *Shigella* survival has not been studied. In addition, *Shigella* are further exposed to plasma during bacteremia, which occurs in most severe cases of shigellosis (2), especially in children under five years of age (3–6) and in immunocompromised or cancer patients (7–9). All species of *Shigella* can cause bacteremia to the same extent (*S. flexneri, S. sonnei, S. boydii* and *S. dysenteriae*) (10). The ability of *Shigella* to reach the bloodstream and cause bacteremia significantly increases the mortality rate in shigellosis patients (11, 12). Patients suffering from shigellosis have been found to have endothelial damage, associated with hemorrhage, congestion and dilation of capillaries in the intestinal mucosa (13).

Therefore, we hypothesized that the ability of *Shigella* to survive exposure to plasma components, both locally during hemorrhage and systemically, contributes to its virulence and may play a central role in shigellosis. The virulence of *Shigella* relies on the expression of two major secretion systems that are conserved among *Shigella* strains: the type III secretion system (T3SS), which is involved in host cell invasion, and the type V secretion system (T5SS). The major T5SS virulence factors secreted by *Shigella* strains belong to the serine protease autotransporter of *Enterobacteriaceae* (SPATE) family. SPATEs consist of two domains: a transmembrane β-barrel domain and a passenger domain, the latter being secreted upon autocleavage within the linker sequence. Two classes (I & II) of SPATEs have been defined according to their structure and properties (14). Like all pathogenic *E. coli* strains, all virulent *Shigella* species secrete at least one member of the SPATE family; they are the most abundant virulence factors released by *Shigella* strains. *S. flexneri* 5a secretes SepA (class II), *S. sonnei* secretes SigA, *S. flexneri* 2a secretes SepA, SigA, and Pic (class II). SepA was the first *Shigella* SPATE to be identified (15).

Local hemorrhage and bacteremia are induced at late stages of *Shigella* infection, which until recently could not be studied in available animal models of shigellosis, because mice are not susceptible to *Shigella* infection and young guinea pigs are only transiently infected by *Shigella* (16). Our team recently validated a new ascorbate-deficient guinea pig as a model for shigellosis which allows to follow prolonged *Shigella* infections (17). In this model, we reported that *S. flexneri* 5a 48h infection induced bacteremia, whereas the molecular mechanisms remained elusive. In this study, we aimed to investigate the contribution of *Shigella* secretion systems in the late phase of infection, especially when *Shigella* is exposed to plasma components during both local hemorrhage and bacteremia.

We have previously shown that *Shigella* forms hypoxic foci of infection during the invasion and colonization of the colonic mucosa, and develops mainly under low-oxygen conditions in the late stages of infection (18), including during bacteremia, as we reported that the blood plasma is poorly oxygenated (pO_2_= 8.4 mmHg) (19). We have previously shown that the T3SS is inactive in the absence of oxygen (18, 20), and we hypothesized that the ability of *Shigella* to secrete virulence factors belonging to the T5SS may be involved in its ability to disseminate in the host at later stages of the infectious process. The contribution of SPATEs to the virulence of *Shigella* is far from fully understood, although some specific targets have been identified. It has been previously reported that Pic degrades O-glycosylated proteins on the surface of leukocytes (21), whereas SepA cleaves neutrophil alpha-1 antitrypsin (22). Pic has previously been reported to cleave complement component C3 *in vitro* (23), suggesting that SPATEs may impair activation of the complement system and maintain the ability of *Shigella* to survive in plasma, although no direct evidence has yet been presented.

In this study, we confirmed that *S. flexneri* 5a and *S. sonnei* induce hemorrhage and reach blood vessels located in the colonic mucosa after prolonged infection. We demonstrated that all the tested *Shigella* strains survive plasma exposure, in contrast to non-pathogenic *E. coli*. We have also shown that the survival of *S. flexneri* 5a and *S. sonnei* depends on the secretion of SepA and SigA, respectively, which both cleave the complement C3, as previously reported for Pic (23). We found that SPATE-dependent cleavage of C3 impairs the formation of the complement membrane attack complex (MAC) (24), and we provided further evidence that SPATEs are essential *in vivo* for the dissemination of *Shigella* in the hemorrhagic colonic mucosa and in the bloodstream during bacteremia, revealing a novel function of SPATEs in the virulence cycle of *Shigella*.

## Results

### *Shigella* spp. induce hemorrhage, reach blood vessels, and survive the plasma exposure

To evaluate the occurrence of the interaction between *Shigella* and plasma components during the infectious process, we first analyzed the infected colonic mucosa of ascorbate-deficient guinea pigs after prolonged infection. We first confirmed that hemorrhage was induced 48h p.i. in abscesses formed by *S. flexneri* 5a and *S. sonnei*, as revealed by the detection of red blood cells (RBCs), which were not observed in uninfected tissues (Fig. 1A). These results are consistent with previous observations made in the human intestine of patients with shigellosis (13). In this princeps study, in addition to hemorrhage, the authors reported endothelial damage associated with shigellosis, possibly associated with peripheral vascular insufficiency. We report here for the first time that *S. flexneri* 5a and *S. sonnei* reached CD31+/CD34+ blood vessels (Fig. 1B). More specifically, most *Shigella* infectious foci detected in the colonic mucosa colocalized with blood vessels that appeared largely disorganized (Fig. 1B). It has been suggested that this ability of *Shigella* to reach blood vessels is related to bacteremia, as reported previously (25). Detection of hemorrhagic abscesses in colonic mucosa infected by *Shigella* and its detection in blood vessels confirmed exposure to plasma components in colonic mucosa.

**Figure 1.**
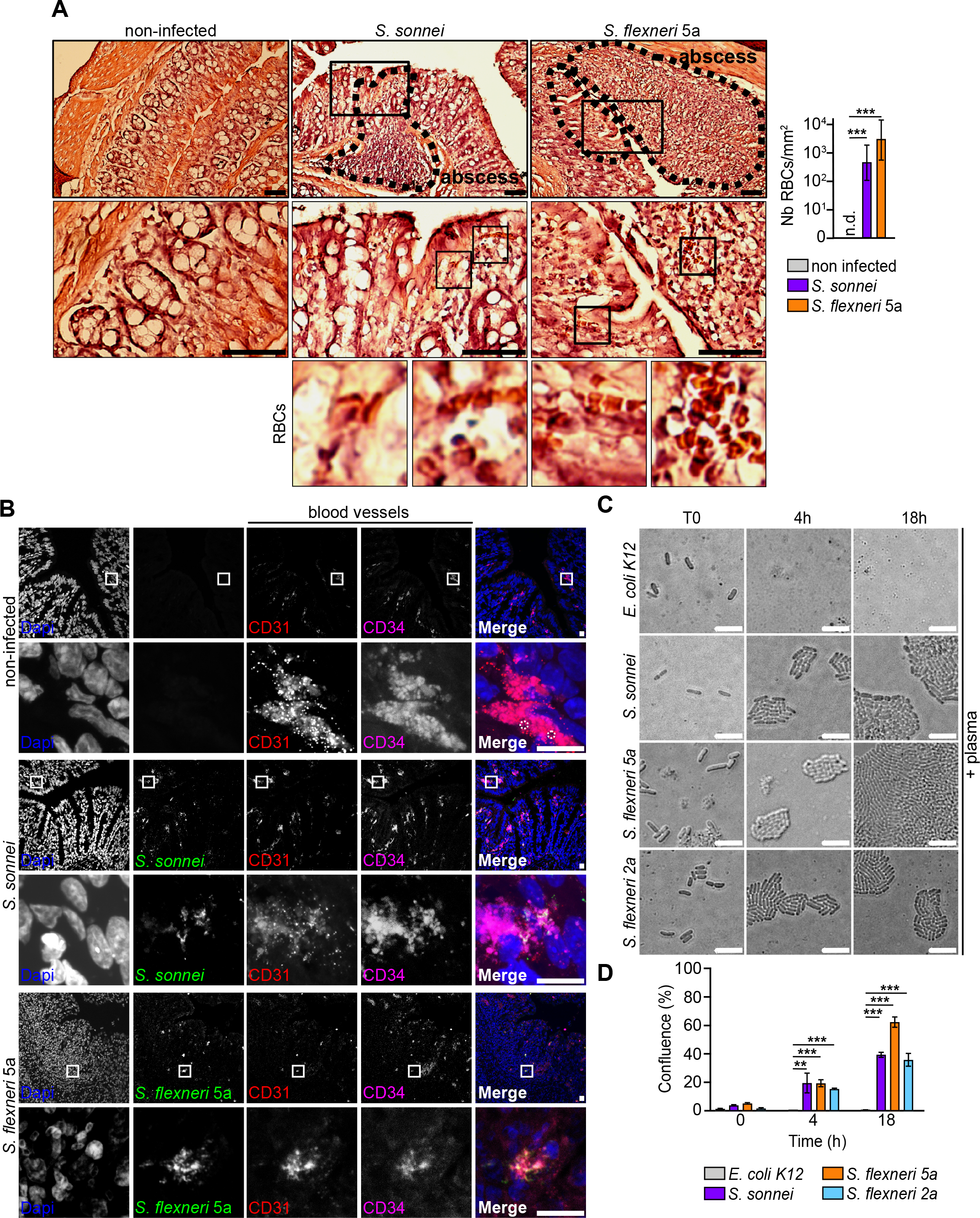
*Shigella* induces local hemorrhage, reaches blood vessels *in vivo* and survives to plasma exposure *in vitro*. **(A)** Guinea pig colonic mucosa infected for 48h with *S. sonnei et S. flexneri* 5a wild-type strains. Black dotted lines delineate abscesses. Bars, 100 μm. Hemorrhage is associated with infiltration of Red Blood Cells (RBCs) within infected tissues, which were counted (Nb RBCs/mm^2^) in each condition. Results are expressed as mean ± S.D (n=3). ‘n.d.’ means not-detected, *** indicates t-test *p* < .001. **(B)** CD31+/CD34+ colonic blood vessels were immunodetected in each condition (red/magenta), together with *S. sonnei et S. flexneri* 5a (green). DNA was stained with Dapi (blue). White boxes indicate individual blood vessels. Capillary lumens are figured with white dashed circles. Bars, 20 μm **(C-D)** Growth of *E. coli* K12, *S. sonnei, S. flexneri* 5a and 2a strains on M9 agar pads in the presence of fresh human plasma at 37°C for 4h and 18h (C). Quantification of bacterial growth (D) by calculating confluence (%) of cultures. Results are expressed as mean ± S.D (n=3). ** indicates t-test *p* < .01, *** indicates t-test *p* < .001.

We further investigated the capacity of *Shigella* to survive and grow in the presence of plasma using M9 agar pad devices, that allowed bacterial growth to be monitored at the single cell level (Fig. S1A). We demonstrated that all tested *Shigella* strains (*S. flexneri* 5a, *S. flexneri* 2a, and *S. sonnei*) were able to survive and grow in the presence of human plasma, in contrast to non-pathogenic *E. coli* K12 which appeared to be lysed at early time points (Fig. 1C-D). As a control, we confirmed that all strains grew at the same rate in the absence of plasma (Fig. S1B-C). When all *Shigella* strains grew in the presence of plasma, we noted a significant reduction in bacterial growth rate in the presence of plasma compared to PBS, particularly at later time points (18h) (Fig. S1D), suggesting that the ability of *Shigella* to survive plasma exposure was associated with an active virulence mechanism; plasma exposure represented a stress to which *Shigella* had to respond to.

In addition, we aimed to identify the molecular mechanisms involved in the survival of *Shigella* upon plasma exposure. We have previously shown that plasma is poorly oxygenated (19) and that *Shigella* develops mainly under low-oxygen conditions within foci of infection (18). Under these conditions, we have shown that *Shigella* T3SS is inactive (20). Alternatively, we hypothesized that *Shigella* SPATEs might be involved in the ability of *Shigella* to survive in plasma, because Pic has previously been reported to target plasma component C3 (23).

### *Shigella* SPATEs: structure and regulation of their expression/secretion

SigA, SepA, and Pic are the only SPATEs secreted by *Shigella* species. We confirmed by mass spectrometry analysis of bacterial cultures that *S. sonnei* secretes only SigA, *S. flexneri* 5a secretes only SepA and *S. flexneri* 2a secretes SigA, SepA and Pic (Fig. 2A). Since only partial information were available on the structure, secretion, and expression regulation of SPATE, we provided additional missing data during this study.

**Figure 2.**
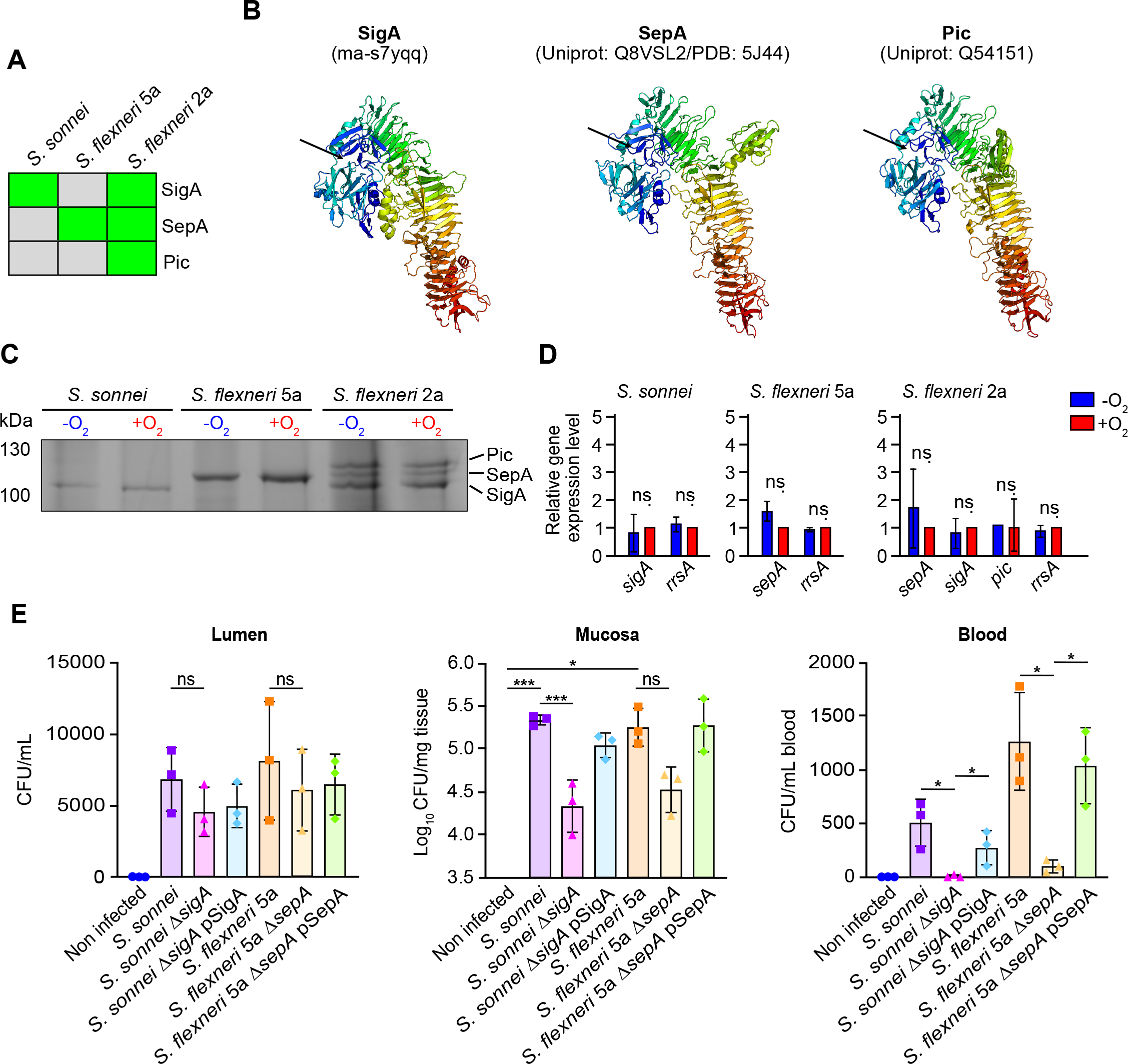
SPATEs structure, secretion regulation and importance during *Shigella* infection. **(A)** SPATEs (SigA, SepA, Pic) secreted by *S. sonnei, S. flexneri* 5a and *S. flexneri* 2a wild-type strains were identified by mass spectrometry. **(B)** The 3D structure of *S. sonnei* SigA was generated with AlphaFold 2 (ModelArchive 10.5452/ma-s7yqq) and compared to the available structures of SepA (Uniprot: Q8VSL2/PDB: 5J44) and Pic (Uniprot: Q54151) (25-27). A rainbow color gradient was used from dark blue (N-terminus) to dark red (C-terminus). Black arrows show position of catalytic triad. **(C)** SPATE-containing culture supernatants from *S. sonnei, S. flexneri* 5a and *S. flexneri* 2a grown in -O_2_ and +O_2_ conditions. Samples were separated on a 10% SDS-PAGE gel and stained with Coomassie. Representative result of 3 independent experiments. **(D)** The expression of SPATE-encoding genes was quantified by qRT–PCR in *S. sonnei, S. flexneri* 5a and *S. flexneri* 2a grown in -O_2_ and +O_2_. *rrsA* mRNA levels were used as a control. Relative gene expression levels are expressed as mean ± S.D (n=3). ‘ns’ indicates t-test *p* > .05. **(E)** The presence of indicated *Shigella* strains (wild-type, SPATE mutants and complemented strains) in the colonic lumen and mucosa and in the blood circulation of guinea pigs was assessed 48h p.i.. Results are expressed as mean ± S.D (n=3). ‘ns’ indicates t-test *p* > .05, * indicates t-test *p* < .05, *** indicates t-test *p* < .001.

The structure of the passenger domain of SepA was previously solved by X-ray crystallography (Q8VSL2 (Uniprot) or 5J44 (PDB)) (26) (Fig. 2B). The structure of the passenger domain of Pic (Q54151 (Uniprot)) (Fig. 2B) was solved using the structure prediction tool Alphafold (27, 28), which allows 3D structure prediction starting from primary sequence information. Since the 3D model of the passenger domain of *S. sonnei* SigA was not available, we obtained its molecular model (Uniprot-ID:Q3YXF8 from residue Met56 to Asn1008) using the structure prediction tool AlphaFold and ColabFold (29), a Web-interfaced implementation of AlphaFold 2. Five models were generated and ranked based on mean pLDDT quality scores, the standard metric of AlphaFold. All five pLDDT scores were above 91 out of a maximum of 100. The top-ranked model with a score of 92.1 was selected for further analysis (Fig. 2B).

To check the validity of the model, we performed a BLASTp similarity search in the Protein Data Bank (PDB) using the SigA passenger domain as a query. The first hit we found was the structure of a homologous protein, with an identity of 46.4% and a similarity of 65%, representing the *E. coli* serine protease EspP (PDB-ID:3SZE and Swissprot-ID:O3259) (30). Our 3D model was superposed on the retrieved 3SZE structure and had a root mean square deviation (RMSD) of 3.6 angstrom (Fig. S2A) showing high similarity between the structures. As expected, the pLDDT scores along the SigA passenger domain were very good, except for local loop regions, which are generally more flexible and/or present less structured elements, making them more difficult to model (Fig. S2B-C). Moreover, the predicted distance errors between protein residues were very low, which supports the quality of the SigA model (Fig. S2D). We further analyzed the structural properties of our AlphaFold model of SigA. A globular N-terminal region that folds into both α-helices and β-strands is connected to an elongated triangular prism of parallel β-strands. Interestingly, the Leu497 -Arg560 region exits the prism and contributes to the folding of the globular region, while the Lys598 to Gly628 region folds to form an antiparallel β-sheet and a loop, both lying on the prism. The catalytic residues (Ser258, His126 and Asp154) are localized in the globular region of the passenger domain, as expected (Fig. S2E).

We further investigated how the secretion of *Shigella* SPATEs is regulated by microenvironmental factors.

It was reported that the secretion of SPATEs was thermoregulated in enteroaggregative *E. coli* (Pic (31)), *S. flexneri* 5a (SepA (15)), and *S. sonnei* (SigA (32)), although we did not observe significant temperature-dependent regulation of SPATE secretion in *S. sonnei, S. flexneri* 5a, and *S. flexneri* 2a (grown at 30°C, 37°C, or 42°C) (Fig. S3A). *Shigella* SPATEs were secreted by all *Shigella* strains tested regardless of temperature. In contrast, the oxygen-dependent regulation of SPATEs expression and secretion has not yet been studied, especially in poorly oxygenated environments such as plasma. We demonstrated that the secretion of SPATEs in *S. flexneri* 2a (SepA, Pic, SigA), *S. flexneri* 5a (SepA), and *S. sonnei* (SigA) was independent of oxygen availability, as observed by Coomassie staining (Fig. 2C) and confirmed by western blot analysis using specific α-SPATE antibodies (Fig. S3B). Consistently, at a transcriptional level, no-oxygen-dependent regulation of the expression of SPATE encoding genes was observed in the tested *Shigella* strains (Fig. 2D). In other words, we demonstrated that SPATEs are produced and secreted by *S. sonnei, S. flexneri* 5a and 2a in the absence of oxygen, a condition encountered by *Shigella* in hypoxic foci of infection and in plasma (19), supporting their potential involvement during hemorrhage or bacteremia. To proceed, *S. flexneri* 5a and *S. sonnei* were used as models because they express and secrete only one member of the SPATE family (SepA and SigA, respectively).

### SPATEs are essential for Shigella dissemination in the blood in vivo

To define the importance of SPATEs during the *S. flexneri* 5a and *S. sonnei* virulence cycles *in vivo, S. sonnei* Δ*sigA* and *S. flexneri* 5a Δ*sepA* mutant strains were constructed, together with complemented strains.

Upon intrarectal challenge of ascorbate-deficient guinea pigs (17), we demonstrated that *S. sonnei* Δ*sigA* and *S. flexneri* 5a Δ*sepA* mutants were attenuated upon prolonged infections (up to 48h) using complementary methods (Fig. 2E and S3C-E). First, we showed that the weight of animals infected by *S. sonnei* Δ*sigA* and *S. flexneri* 5a Δ*sepA* strains was significantly higher 48h p.i. compared to animals infected by the corresponding wild-type or complemented strains, with marked weight gain between 24h p.i. and 48h p.i. (Fig. S3C). Since *Shigella* infection severity is correlated to guinea pig weight loss, these results were consistent with the reduced virulence of *S. sonnei* Δ*sigA* and *S. flexneri* 5a Δ*sepA* mutant strains. To correlate these results with the bacterial load in the colon and in the bloodstream, CFU counts were assessed in lumen, mucosa and in blood (Fig. 2E).

CFU counting revealed that *S. sonnei* Δ*sigA* and *S. flexneri* 5a Δ*sepA* mutant strains had a reduced ability to colonize the colonic mucosa, and unable to survive in animal blood, in contrast to the wild-type and complemented strains (Fig. 2E). We confirmed the absence of RBCs (hemorrhage markers) in the colonic mucosa of guinea pigs infected by *S. sonnei* Δ*sigA* and *S. flexneri* 5a Δ*sepA* mutant strains (Fig. S3D), as observed with wild-type strains (Fig. 1A), suggesting that these mutant strains did not alter the colonic mucosa microvasculature. We confirmed that *S. sonnei* Δ*sigA* and *S. flexneri* 5a Δ*sepA* mutant strains were not detected in CD31+/CD34+ blood vessels (Fig. S3E), in contrary to wild-type strains (Fig. 1B).

These results strongly suggested that SPATEs contribute to the survival of *Shigella* strains to plasma exposure, which occurs in hemorrhagic tissues and in the bloodstream.

### SPATEs promote the survival of *Shigella* to plasma exposure

We demonstrated that *S. sonnei* Δ*sigA* and *S. flexneri* 5a Δ*sepA* mutant strains treated with plasma at all time points had a significant growth defect (Fig. 3A-B), and we confirmed in the absence of plasma that both mutant strains had no growth defect (Fig. S4A), confirming the importance of SPATEs in the ability of *Shigella* strains to survive in plasma. We showed by live microscopy the growth defect due to plasma of both *S. sonnei* Δ*sigA* and *S. flexneri* 5a *ΔsepA* (Fig. S4B-C). The growth defect of the *S. flexneri* 5a Δ*sepA* mutant strain in the presence of plasma, although significative, was weaker as compared to *S. sonnei* Δ*sigA* mutant strain, compared to corresponding wild-type strains (Fig. 3A-B), we further aimed at confirming the importance of SepA in the capacity of *S. flexneri* 5a to survive and grow in the presence of plasma. Since the significant growth defect of the *S. flexneri* 5a Δ*sepA* mutant was less pronounced than that of *S. sonnei* Δ*sigA* mutant, we sought to address the importance of SepA in the growth and survival of *S. flexneri* 5a in plasma. We demonstrated that the BS176 strain (Plasmid-cured *S. flexneri* 5a M90T) had a growth defect in the presence of plasma and had a rounded-cell phenotype (Fig. S4D). We showed that the overexpression of SepA in BS176 (BS176 pSepA strain) suppressed the growth defect in the presence of plasma, but not the overexpression of the catalytically inactive SepA_S211A_ (BS176 pSepA_S211A_ strain) (Fig. S4D). These complementary results confirmed the contribution of SepA to the survival of *S. flexneri* 5a in plasma.

**Figure 3.**
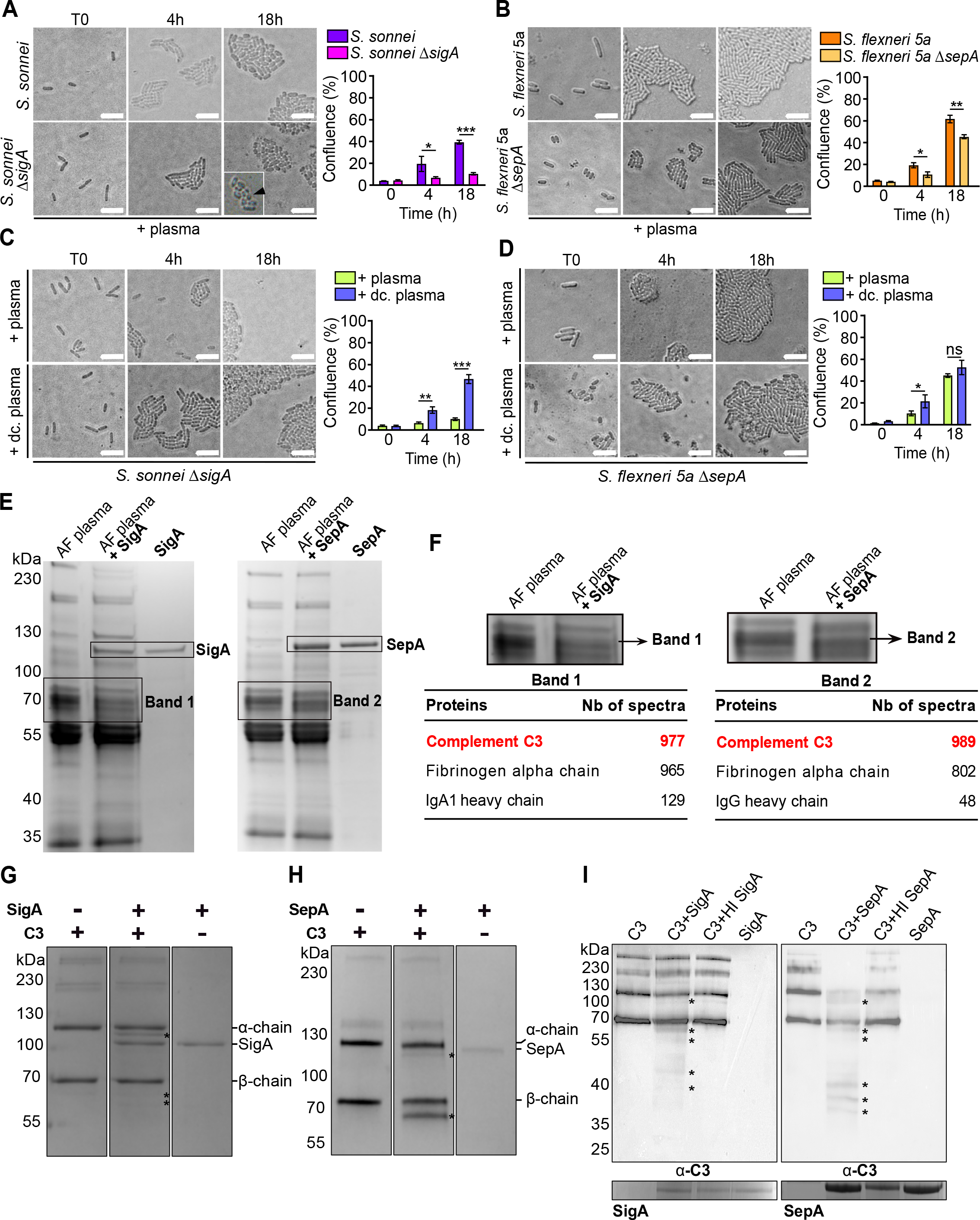
SPATEs are essential for *Shigella* survival to plasma exposure, by cleaving Complement C3. **(A-B)** *S. sonnei* wild-type and Δ*sigA* strains (A) and *S. flexneri* 5a wild-type and Δ*sepA* strains (B) were incubated up to 18h in presence of fresh human plasma on M9 agar pads and the bacterial confluence was quantified. Results are expressed as mean ± S.D (n=3). ‘ns’ indicates t-test *p* > .05, * indicates t-test *p* < .05, ** indicates t-test *p* < .01, *** indicates t-test *p* < .001. **(C-D)** *S. sonnei* Δ*sigA* (C) and *S. flexneri* 5a Δ*sepA* (D) strains were incubated in the presence of fresh human plasma or decomplemented plasma (dc. plasma) and data were analyzed as in (A-B). **(E)** Human albumin-free plasma (AF plasma) was incubated O/N at 37 °C with purified SigA or SepA and was analyzed on 10% SDS-PAGE gel, stained with Coomassie. **(F)** Mass spectrometry analysis of Band 1 and 2, as indicated (from gel in (E)). **(G-H)** Purified human complement component 3 (C3, α-chain/β-chain) was incubated with purified SigA and SepA for 18 hours at 37°C, samples were processed as in (E). **(I)** Impact of SigA and SepA heat-inactivation (HI) on C3 cleavage. Samples were analyzed by western blot using human anti-C3 antibody (top) and SigA and SepA were stained with Coomassie (bottom).

Since it was previously reported that Pic targeted C3 (23), we hypothesized that the activation of the plasma complement system was responsible of the growth defect of *S. sonnei* Δ*sigA* and *S. flexneri* 5a Δ*sepA* mutant strains. Indeed, we demonstrated that the growth defect in plasma was reversed when plasma was decomplemented(Fig. 3C-D) with the mutants growing at the same rate as wild-type strains (Fig. S4E-F). These results supported our hypothesis and we further aimed to identify the complement component, which may be targeted by SigA and SepA.

### SigA and SepA cleave complement component 3 (C3)

To identify SPATE targets in plasma, we used an unbiased approach consisting of incubating albumin-free plasma with purified SigA or SepA (Fig. 3E). Removing albumin, the major protein in plasma, was mandatory to visualize less abundant plasma proteins at the same molecular weight (67 kDa). In the presence of SigA or SepA, plasma proteins located in an identified band around 70 kDa disappeared (Fig. 3E). By mass spectrometry analysis, we identified complement component 3 (C3) as the most abundant protein composing both bands (1 and 2) (Fig. 3F). It must be noticed that fibrinogen alpha chain was also abundant in these bands (Fig. 3F). Since to our knowledge, fibrinogen had no direct antimicrobial activity, we focused on the cleavage of C3 by SPATEs. Commercially purified C3 was incubated with purified SigA and SepA and Coomassie staining revealed new bands confirming SPATEs target C3 (Fig. 3G-H). We further identified C3-degradation products by western blotting, and we showed that heat-inactivated SigA and SepA were no longer able to cleave C3 (Fig. 3I).(As controls,) we further demonstrated that SigA and SepA did not cleave complement C5 or various immunoglobulins such as IgG, IgM, IgA (Fig. S5A-B). To identify the SPATE cleavage site on C3, the sequences of C3 α-chain, β-chain and the cleaved α-chain formed in the presence of SigA were analyzed by mass spectrometry. We could only identify one potential cleavage site of SigA, located in the complement C3 α-chain between residues 680 and 740 (Fig. S5C). Despite several attempts, we failed to proceed similarly with C3 incubated with SepA (data not shown).

Taken together, our results showed that SPATEs contribute to the impairment of the complement system by *Shigella*, allowing its survival to plasma. Since we demonstrated that plasma content was released within the infected colonic mucosa, we further aimed to confirm the presence of C3 and other plasma components such as albumin, the most abundant plasma protein, in this microenvironment.

### *Shigella-*C3 interaction occurs within hemorrhagic infected tissues

Since we demonstrated that hemorrhage was caused by *Shigella* infection with the detection of RBCs in the colonic mucosa (Fig. 1A), we further aimed to confirm this result and evaluate the distribution of plasma components (C3 and albumin) in relation with *Shigella* dissemination. In non-infected tissues, we could first immunodetect C3 and albumin strictly localized within blood vessels (Fig. 4A-B). We observed that C3 and albumin were more abundant in the colonic mucosa infected by *S. sonnei* and *S. flexneri* wild-type strains compared to the uninfected animals, and we demonstrated that *Shigella* foci co-localized with both C3 and albumin, confirming the interaction between *Shigella* and plasma components in hemorrhagic tissues (Fig. 4A-B). As control, we did not observe changes in C3 and albumin abundance and distribution with tissues infected by *S. sonnei* Δ*sigA* and *S. flexneri* 5a Δ*sepA* mutant strains compared to the non-infected tissues (Fig. S6A-B), confirming the absence of hemorrhage in this context, as previously reported (Fig. S3D-E).

**Figure 4.**
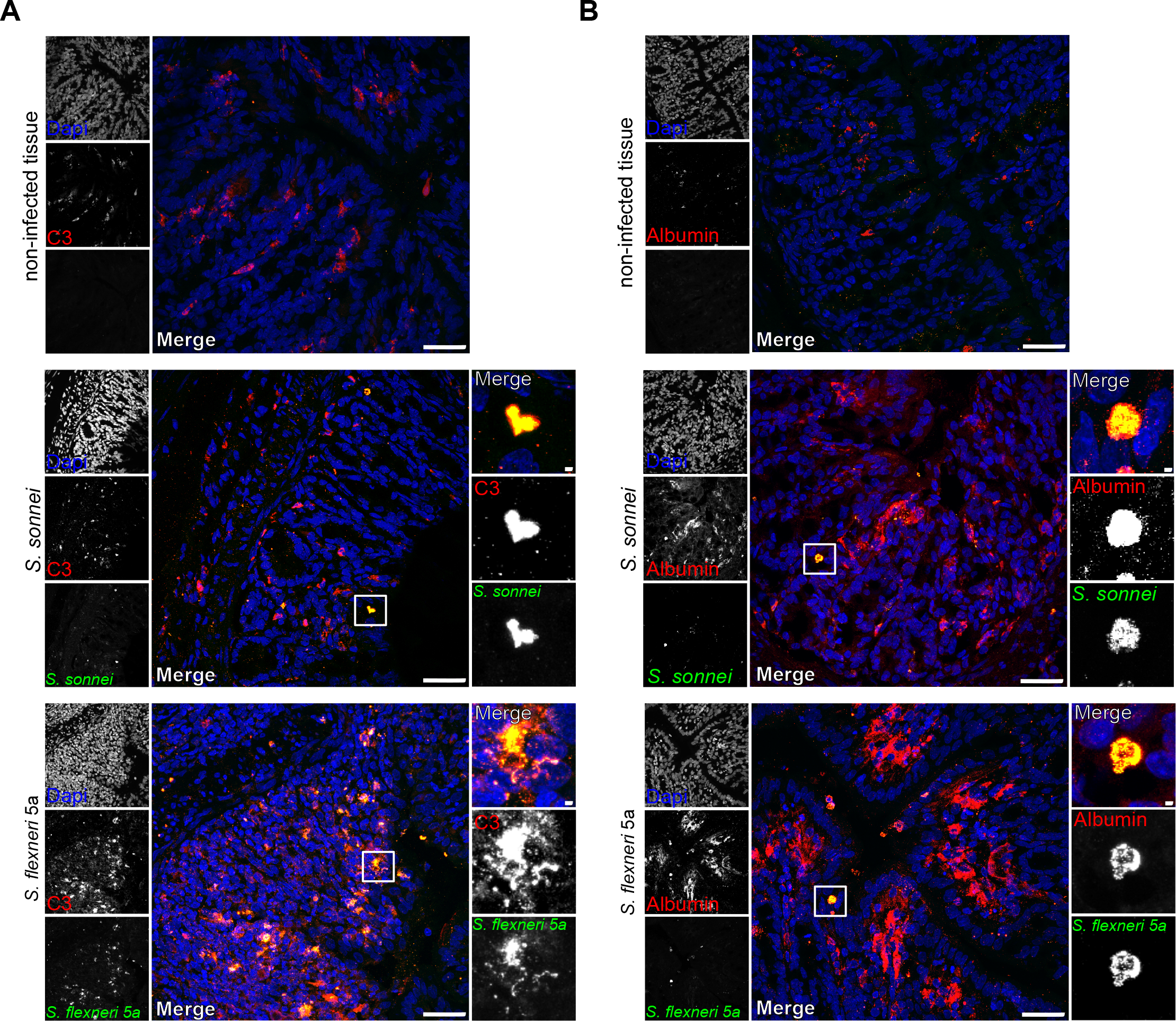
*Shigella* are exposed to complement C3 *in vivo* during hemorrhage. **(A)** Guinea pig colonic mucosa infected by *S. sonnei* and *S. flexneri* 5a wild-type strains (green) for 48h. Infected and non-infected tissues were stained with an α-human complement C3 antibody (red) and DNA was stained with Dapi. Bars, 50 μm. **(B)** Tissue hemorrhage was further emphasized in using an α-human albumin antibody (red) upon infection by *Shigella* (green). DNA was stained with Dapi. Bars, 50 μm.

## Discussion

In this report, we demonstrated that all tested *Shigella* strains survive exposure to plasma (Fig. 1C) which not only occurred during bacteremia but also upon hemorrhage induced by *Shigella* in the colonic mucosa (Fig. 1A). We demonstrated that SPATEs contribute significantly to the ability of *Shigella* to survive plasma exposure by cleaving the central component of the complement system, the complement component 3 (C3) (Fig. 3E-I). Our results shed new light on the antimicrobial activity of the complement system, which is active outside of the blood circulation, within hemorrhagic tissues, like the colonic mucosa infected by *Shigella*.

The invasive attenuation of *S. sonnei* Δ*sigA* and *S. flexneri* 5a Δ*sepA* mutant strains (Fig. 2E, S3C-E and S6A-B) may be explained by their inability to resist the complement system within the colonic mucosa, hence impairing their dissemination within infected tissues. It must be noticed that additional virulence mechanisms involved in the complement system subversion may be expressed by *S. flexneri* 5a, because the susceptibility to plasma of the *S. flexneri* 5a Δ*sepA* mutant strain was less pronounced than the *S. sonnei* Δ*sigA* mutant strain (Fig. 3A-B).

Additionally, in this study, we characterized molecular aspects of the hemorrhage induction during shigellosis, thanks to the validation of a new animal model of shigellosis which allowed us to follow prolonged infection episodes (17). We also reported how *Shigella* reached specifically blood vessels in the colonic mucosa at late infection stage (Fig. 1B). Since we previously demonstrated that *Shigella* aerobic respiration was essential to the formation of hypoxic foci of infection, further investigations will be required to better appreciate the importance of endothelial damages caused by *Shigella*, and previously reported in shigellosis patients (13), in the maintenance of low-oxygen levels within infected tissues and its importance to the development of the disease.

## Materials and Methods

### Bacterial strains and plasmids

All strains and plasmids used in this study are listed in Table S1. *S. sonnei* wild-type strain was obtained from the Pasteur Institute collection (CIP 106347). *S. flexneri* 2a and *S. flexneri* 5a wild-type strains belong to the laboratory collection. BS176 is *S. flexneri* 5a cured from its pINV virulence plasmid (33). *E. coli* K12 (*Escherichia coli* str. K-12 substr. MG1655) was used as a control. *S. sonnei* Δ*sigA* and *S. flexneri* 5a Δ*sepA* mutant strains were constructed with the lambda-red recombination method (34). In brief, *sigA* and *sepA* genes were replaced by a chloramphenicol resistance cassette, which was amplified from the PKD3 plasmid using primers listed in Table S2. Complementation of *S. sonnei* Δ*sigA* and *S. flexneri* 5a Δ*sepA* mutant strains was performed with the plasmid pSigA3 (pSigA, AmpR) (kindly provided by M. Meza-Segura (35)) and pZK15 (pSepA) (36),(kindly provided by Pr. Claude Parsot) respectively. The strain *E. coli* HB101 pPic1 (37) was used to overexpress and purify the protein Pic. *Shigella* strains were grown in TSB liquid medium and on TSB petri dishes supplemented with 0,1% Congo Red and *E. coli* K12 strain was grown in LB liquid medium and LB agar plates. Antibiotics were added when required at the following concentrations: Ampicillin 100μg/ml, Chloramphenicol 30 μg/ml, Tetracycline 10μg/ml.

### SigA, SepA and Pic purification

SigA was purified from *S. sonnei* culture supernatant. Briefly, bacteria were grown overnight at 37°C and subcultured in a fresh medium for 5 h at 37°C. SepA and Pic were purified from BS176 pSepA and *E. coli* HB101 pPic strain, grown overnight at 37°C.

#### SigA purification

The bacterial culture was centrifuged for 15 min at 4.000 x *g* at 4°C and the supernatant was filtered through a 0.22μm filter and subsequently precipitated in ammonium sulfate (35% w/v) for 45 min at 4°C under gentle agitation. The precipitated supernatant was centrifuged for 30 min at 7.500 x *g*, and the pellet was resuspended in 25mM NaH_2_PO_4_, 25mM Na2HPO_4_, 50mM NaCl buffer (pH 7.5) and dialyzed overnight in the same buffer. The dialyzed sample was loaded onto a DEAE column (TSK-gel DEAE-5PW - Tosoh) and a NaH_2_PO_4_ 25mM/Na2HPO_4_ 25mM/NaCl 1M gradient (pH 7.5) was applied. Purified SigA was collected, quantified, and stored at 4°C (for up to one month).

#### SepA and Pic purification

Bacterial cultures were centrifuged for 15 min at 4000 x *g* at 4°C and the supernatant was filtered through 0.22μm filters and subsequently precipitated in ammonium sulfate (30% w/v) for 45 min at 4°C under gentle agitation. Precipitated samples were centrifuged for 30 min at 7500 x *g* at 4°C and pellets were resuspended in 1X Phosphate Buffer Saline (PBS). Samples were concentrated on a 100kDa cut-off Amicon filter (Merck) and loaded on a gel filtration column (Superdex 200 10/300 GL – GE Healthcare). Purified SepA and Pic were collected, quantified, and stored at -20°C (for up to one month).

### Production of anti-SPATEs antibodies

Rabbit polyclonal antibodies specific of SigA, Pic and SepA were produced. Briefly, SigA, Pic, and SepA were purified by ammonium sulfate precipitation, concentrated, and further dialyzed. New Zealand rabbits were immunized by the Genecust company through series of injections. Antibodies were purified on by affinity chromatography. Antibodies were stored at -20°C in PBS supplemented with 0,02% NaN_3_. Antibodies were further purified by affinity to the corresponding SPATE transferred on a nitrocellulose membrane from supernatants of *S. flexneri* 5a BS176 pSepA, *S. sonnei* Δ*sigA* pSigA and *E. coli* HB101 pPic, after a migration SDS-PAGE. Specific bands corresponding to SepA, SigA or Pic were cut, blocked with PBS supplemented with 0.1% Triton X-100 + 10% Fetal Bovine Serum and incubated overnight with anti-SepA serum, anti-SigA serum or anti-Pic serum respectively. After several washes with the blocking buffer, bound antibodies were collected using Glycine 0.2M pH2.8 supplemented with 0.2% Gelatin, then the pH was quickly neutralized with 25% v/v unbuffered Tris 1M. Purified antibodies were tested on whole supernatants to assess their purity.

### Human plasma collection and treatment

Human blood was collected from the antecubital vein of anonymous voluntary donors at the Etablissement Français du Sang de Strasbourg (authorization n°ALC/PIL/DIR/AJR/FO/606). Blood bags of 500 mL were collected and rapidly transferred into an anoxic chamber (DG250, Don Whitley). The platelet-free plasma fraction was collected by centrifugation for 20 min at 3.800 rpm and was further stored at -20°C. Decomplemented plasma was obtained by heating plasma for 30 min at 56°C and was further stored at -20°C. Albumin-free plasma was obtained following the Minute^™^ Albumin Depletion Reagent for Plasma and Serum kit instructions (Invent Biotechnologies) and was stored at -20°C.

### Bacterial growth assay

Bacteria were grown in M9 minimal medium (Merck) supplemented with 2 mM MgSO4, 0,1 mM CaCl_2_, and 0,4% glucose overnight at 37°C (180 rpm). Bacterial cultures were diluted to OD_600nm_0.05 in fresh rich M9 media and were grown for 2 hours at 37°C to reach OD_600nm_ 0,22. 1μl of bacterial culture and 5μl of PBS, plasma or decomplemented plasma were deposited onto a 1% M9-agar pad. Pads were then incubated for up to 18 hours at 37°C in humid conditions to allow bacterial growth. Images were taken at T=0, T=4h, and T=18h with an Axioskop 2 light microscope (Zeiss, Germany) equipped with an Optikam Pro6 digital Camera (Optika, Italy) and a X100 objective (Plan-NEOFLUAR 100X 1.3NA Ph3 oil). Bacterial growth quantification was assessed by quantifying the culture confluence (expressed as %), using the Fiji v2.1.0 software.

### SigA structure prediction

To predict the structure of the passenger domain of *S. sonnei* SigA, the protein sequence Uniprot-ID:Q3YXF8 (NCBI protein sequence WP_052993189) from residue Met56 to Asn1008 was submitted to Colabfold v1.5.2 that implements AlphaFold2. Five models were requested with AMBER force field relaxation. The BLASTp similarity search with SigA passenger domain as a query was carried out with default parameters (Expect threshold 0.05, word size 5, BLOSUM62 scoring matrix, gap creation penalty 11, gap extension penalty 1) in the PDB. Structure manipulations were carried out in PyMOL 1.8.4. The align function which is based on primary sequence comparison was used to superimpose structures. Calculations of RMSD were run in Visual molecular dynamics (VMD) 1.9.3. The predicted structure of SigA was deposited with ModelArchive and is publicly available (https://modelarchive.org/doi/10.5452/ma-s7yqq).

### SPATEs secretion regulation

To study the regulation of SPATE secretion, bacterial cultures of *S. sonnei, S. flexneri* 2a and *S. flexneri* 5a were grown O/N at 37°C (180 rpm). The following day, cultures were diluted in fresh culture medium at an initial OD_600nm_ of 0.05. To study the O_2_-dependent regulation of SPATE secretion, cultures were incubated at 37°C either under atmospheric conditions with agitation, or in in an anoxic chamber (Don Whitley ; DG250). To study the temperature-dependent regulation of SPATE secretion, cultures were incubated at 30°C, 37°C, or 42°C for 6 hours with agitation (180 rpm) under atmospheric conditions.

In all tested conditions, supernatants were collected by centrifugation for 30 min at 4.000 rpm, and were subsequently filtered (0,22μm) and precipitated in 35% ammonium sulfate (w/v). Precipitated proteins were pelleted by centrifugation (30 min at 7.500 *x g* at 4°C) and further resuspended in 1X PBS. Protein concentration was adjusted by concentrating of the samples on 100kDa filters (Amicon, Merck). Samples were loaded on 10% SDS-PAGE gel and SPATEs were visualized by Coomassie Blue staining (Instant Blue, Abcam) or western blot using specific antibodies. In more details, proteins were transferred on PVDF membranes and rabbit polyclonal antibodies were used at a 1:500 dilution (anti-Pic and anti-SepA) or a 1:250 dilution (anti-SigA) in 1X TBS supplemented with 0,5% of Tween 20 (Merck) and 3% BSA (Euromedex). A horseradish peroxidase-conjugated goat anti-rabbit IgG antibody (Abcam) was used at a 1:10000 dilution and combined to a Super Signal ECL kit (Thermoscientific). Membranes were imaged with a ChemiDoc™ Touch Imaging System (Bio-Rad).

### Quantitative Real-time PCR analysis

*S. flexneri* 2a, *S. flexneri* 5a, and *S. sonnei* were cultured for 5 hours at 37°C (180 rpm) and bacteria were pelleted (15 min centrifugation at 4000 rpm) to proceed with RNA extraction. Total RNAs were extracted by phenol/chloroform and precipitated. After DNAse treatment, RNAs were reverse transcribed using *iScript*^*™*^ *Reverse Transcription Supermix* (Bio-Rad) and cDNAs were amplified by quantitative RT-PCR (qRT-PCR). Reactions were carried out on a CFX96 Real-Time PCR detection system (Bio-Rad) using the Maxima SYBR Green kit (Thermo Scientific). Oligonucleotides used for qRT-PCR are listed in Supplementary Table S2. qRT-PCR reactions were performed and designed according to the MIQE guidelines (38) The specificity of the oligonucleotides was validated, and the amplification efficiencies of the primer sets were between 90 and 110% with r2 values greater than 0.98. Relative mRNA expression levels were calculated using the ΔΔCt method. Results were expressed as mean +/-standard error of an average of three measurements.

### SPATE proteolytic assays

To identify targets, purified SPATEs were incubated with various substrates in 1X PBS at 37°C. Albumin-free plasma and commercially available purified C3, C5, IgG, IgA or IgM (Merck) were incubated for 18 hours at 37°C in the absence or presence of purified SepA or SigA, at the indicated concentrations. Degradation products were separated on 10% SDS-PAGE gels and stained with Instant Blue solution or analyzed by western blot. When indicated, heat-inactivated (30min, 95°C) SigA and SepA were used as controls.

For western blot, O/N degradation products were loaded on a 10% SDS-PAGE gel and then transferred to a PVDF membrane. After 1 hour of blocking, the membrane was incubated O/N at 4°C with gentle agitation with 1:2000 goat anti-human complement C3 antibody (Thermo Scientific) diluted in 1X TBS/0,5% Tween/3% BSA. The next day, the membrane was washed three times in 1X TBS/0,5% Tween and incubated for 1 hour with 1:10 000 horseradish peroxidase-conjugated rabbit anti-goat IgG (Abcam). After three washes, the membrane was revealed using Super Signal ECL kit.

### Mass-spectrometry analysis by nanoLC-MS/MS: in-gel samples, database search and protein validation

For LC-MS/MS analyses of the in-gel samples, samples were loaded on a 4-15% SDS-PAGE precast gel (Bio-Rad) and stain with Coomassie. Bands were excised from the SDS-PAGE gel, were destained several times in 50 mM ammonium bicarbonate containing 50% (v/v) acetonitrile, further dehydrated with 100% acetonitrile and then reduced with 10mM DTT for 1 hour at 56°C. Proteins were then alkylated with 55mM iodoacetamide for 1 hour in the dark at room temperature. Gel pieces were washed again with the destaining solution described above. 80 ng of modified sequencing-grade trypsin (10ng/μL; Promega, Fitchburg, MA, USA) were added to each dehydrated gel sample for overnight digestion at 37°C. Resulting peptides were extracted twice from the gel pieces with a first solution of 60% acetonitrile and 5% formic acid, followed by a second extraction in 100% acetonitrile. Gel supernatants were finally vacuum-dried in a SpeedVac concentrator. Digested peptides were resuspended in 20μL of 0.1% formic acid (solvent A) and injected on an Easy-nanoLC-1000 system coupled to a Q-Exactive Plus mass spectrometer (Thermo Fisher Scientific, Germany). Each sample was loaded on a C-18 precolumn (75 μm ID × 20 mm nanoViper, 3μm Acclaim PepMap; Thermo-Fisher Scientific) and separated on an analytical C18 analytical column (75 μm ID × 25 cm nanoViper, 3μm Acclaim PepMap) with a 60 minutes gradient of solvent B (0.1% of formic acid in acetonitrile).

MS data were searched with Mascot algorithm (version 2.8, Matrix Science) against either the UniprotKB database with *Shigella sonnei* taxonomy (22,219 sequences, version 2021_02), and the swissprot database with human taxonomy (20,396 sequences, version 2021_01) or the swissprot with all taxonomies (568,744 sequences, version 2022_05). The resulting Mascot files were imported into Proline v1.0 package (39) to align the identified proteins. Proteins were then validated on Mascot rank equal to 1, a Mascot score superior to 25, and 1% FDR on both peptide spectrum matches (PSM) and protein sets (based on Mascot score). For figure S5C, the number of observations of each amino acid from the sequences was counted in all the spectra validated with 1% FDR.

### The ascorbate-deficient guinea pig model of shigellosis

The ascorbate-deficient guinea pig model shigellosis was previously described by Skerniskyte et *al*. (17) and was used here to study late infection stages (up to 48h) using the following strains: *S. sonnei* wild-type pGFP, *S. sonnei* Δ*sigA* pGFP, *S. flexneri* 5a wild-type pGFP, *S. flexneri* 5a Δ*sepA* pGFP, *S. sonnei* Δ*sigA* pSigA3, and *S. flexneri* 5a Δ*sepA* pSepA strains. Before the challenge, *sigA* and *sepA* expression was induced with 500μM IPTG from OD_600nm_ 0.3 to 0.6 in *S. sonnei* Δ*sigA* pSigA3 and *S. flexneri* 5a Δ*sepA* pSepA respectively. Briefly, 5-week ascorbate-deficient guinea pigs were intrarectally challenged with 10^11 CFU in the exponential phase (OD_600nm_ = 0.6). Upon infection, animals were weighted on a daily basis and 48 hours post-infection, animals were sacrificed, blood samples were collected by intracardiac puncture and colons were collected and flushed with 500μL PBS. Lumenal CFU counting was performed by plating series of dilutions. Mucosal CFU counting was achieved by plating series of dilutions of homogenized colonic mucosa (0.5 cm tissue sections, weighted and processed in 500μL 1X PBS with Bead Mill Homogenizer, VWR). Blood CFU counting was performed by plating 50μL of blood collected in the presence of an anticoagulant (citrate). For immunofluorescence imaging, 1 cm sections of colons were fixed in PFA, washed with a sucrose gradient (from 15% to 30% sucrose), and frozen in OCT blocks. Then 10μm thickness sections of colons were cut using a CM-3050 cryostat (Leica Biosystems).

### Tissue labeling and imaging

For histological studies, sectioned tissues were stained with hematoxylin-eosin standard protocol and imaged with an Axioskop 2 transmission light microscope (Zeiss) using x10 or x20 objectives. For immunofluorescence, tissues were washed 3 times in PBS/0,1% Tritton, PBS, water, and stained O/N at 4°C in humid conditions with primary antibodies. Following antibodies were used as primary antibodies: 1:100 mouse anti-human CD31 (BD Pharmingen), 1:100 a mouse anti-human CD34 (Biolegend), 1:2000 goat anti-human complement C3 antibody (Thermo Fischer Scientific), 1:1000 goat anti-Albumin antibody (Sigma) and 1:1000 goat anti-Fibrinogen antibody (Thermo Fischer Scientific). The next day, slides were washed three times, and incubated for 1 hour at room temperature with 1:1000 of donkey anti-goat IgG Alexa Fluor 568 (Invitrogen) and DAPI (Life Technologies). Slides were washed again, mounted with ProLongGold® (Invitrogen) and imaged using a laser-scanning TCS SP8 confocal microscope (Leica).

## Acknowledgments

We thank Dr. Annick Dejaegere (IGBMC-Illkirch, France) for supporting the structural analysis of SPATEs. We thank Dr. Mario Meza-Segura for sharing the pSigA plasmid and Dr. Fernando Luiz Perez for sharing the pPic plasmid

## References

1. K. L. Kotloff, M. S. Riddle, J. A. Platts-Mills, P. Pavlinac, A. K. M. Zaidi, Shigellosis. Lancet 391, 801–812 (2018).

2. S. K. Niyogi, Shigellosis. J. Microbiol. (Seoul, Korea) 43, 133–43 (2005).

3. M. J. Struelens, et al., Shigella Septicemia: Prevalence, Presentation, Risk Factors, and Outcome. J. Infect. Dis. 152, 784–790 (1985).

4. C. Nayyar, P. Thakur, V. Tak, A. Singh, Shigella sonnei Sepsis in an Infant: A Case Report. J. Clin. Diagn. Res. 11, DD01–DD02 (2017).

5. M. Delgado, et al., Bacteriemia por Shigella flexneri en dos lactantes. Rev. Chil. infectologa 35, 317–320 (2018).

6. R. M. Kligler, P. D. Hoeprich, Shigellemia. West. J. Med. 141, 375–8 (1984).

7. S. M. Qadri, S. H. Khalil, Polymicrobial septicemia due to Shigella flexneri and pseudomonas aeruginosa: first report. J. Natl. Méd. Assoc. 79, 1289–92 (1987).

8. C.-Y. Liu, Y.-T. Huang, C.-H. Liao, S.-C. Chang, P.-R. Hsueh, Rapidly Fatal Bacteremia Caused by Shigella sonnei without Preceding Gastrointestinal Symptoms in an Adult Patient with Lung Cancer. Clin. Infect. Dis. 48, 1635–1636 (2009).

9. M. Tobin-D’Angelo, et al., Shigella Bacteremia, Georgia, USA, 2002–2012 - Volume 26, Number 1—January 2020 - Emerging Infectious Diseases journal - CDC. Emerg. Infect. Dis. 26, 122–124 (2020).

10. W. A. Khan, J. K. Griffiths, M. L. Bennish, Gastrointestinal and Extra-Intestinal Manifestations of Childhood Shigellosis in a Region Where All Four Species of Shigella Are Endemic. PLoS ONE 8, e64097 (2013).

11. M. J. Struelens, et al., Shigella Septicemia: Prevalence, Presentation, Risk Factors, and Outcome. J. Infect. Dis. 152, 784–790 (1985).

12. J. M. van den Broek, et al., Risk factors for mortality due to shigellosis: a case-control study among severely-malnourished children in Bangladesh. J. Heal., Popul., Nutr. 23, 259–65 (2005).

13. R. Koshi, G. Chandy, M. Mathan, V. I. Mathan, Vascular changes in duodenal mucosa in shigellosis and cholera. Clin. Anat. 16, 317–327 (2003).

14. F. Navarro-Garcia, Serine proteases autotransporter of Enterobacteriaceae: Structures, subdomains, motifs, functions, and targets. Mol. Microbiol. 120, 178–193 (2023).

15. Z. Benjelloun-Touimi, P. J. Sansonetti, C. Parsot, SepA, the major extracellular protein of Shigella flexneri: autonomous secretion and involvement in tissue invasion. Mol. Microbiol. 17, 123–135 (1995).

16. D.-H. Shim, et al., New Animal Model of Shigellosis in the Guinea Pig: Its Usefulness for Protective Efficacy Studies. J Immunol 178, 2476–2482 (2007).

17. J. Skerniskyte, et al., Ascorbate deficiency increases progression of shigellosis in guinea pigs and mice infection models. Gut Microbes 15, 2271597 (2023).

18. J.-Y. Tinevez, et al., Shigella-mediated oxygen depletion is essential for intestinal mucosa colonization. Nat. Microbiol. 4, 2001–2009 (2019).

19. L. Injarabian, et al., Ascorbate maintains a low plasma oxygen level. Sci. Rep. 10, 10659 (2020).

20. B. Marteyn, et al., Modulation of Shigella virulence in response to available oxygen in vivo. Nature 465, 355–358 (2010).

21. F. Ruiz-Perez, et al., Serine protease autotransporters from Shigella flexneri and pathogenic Escherichia coli target a broad range of leukocyte glycoproteins. Proc. Natl. Acad. Sci. 108, 12881–12886 (2011).

22. F. Ruiz-Perez, et al., Serine protease autotransporters from Shigella flexneri and pathogenic Escherichia coli target a broad range of leukocyte glycoproteins. Proc. Natl. Acad. Sci. 108, 12881–12886 (2011).

23. A. G. Abreu, et al., The Serine Protease Pic From Enteroaggregative Escherichia coli Mediates Immune Evasion by the Direct Cleavage of Complement Proteins. J. Infect. Dis. 212, 106–115 (2015).

24. C. B. Xie, D. Jane-Wit, J. S. Pober, Complement Membrane Attack Complex New Roles, Mechanisms of Action, and Therapeutic Targets. Am. J. Pathol. 190, 1138–1150 (2020).

25. A. C. André, et al., The ascorbate-deficient guinea pig model of shigellosis allows the study of the entire Shigella life cycle. bioRxiv, 2020.08.28.270074 (2020).

26. A. Maldonado-Contreras, et al., Shigella depends on SepA to destabilize the intestinal epithelial integrity via cofilin activation. Gut Microbes 8, 544–560 (2017).

27. J. Jumper, et al., Highly accurate protein structure prediction with AlphaFold. Nature 596, 583–589 (2021).

28. M. Varadi, et al., AlphaFold Protein Structure Database: massively expanding the structural coverage of protein-sequence space with high-accuracy models. Nucleic Acids Res. 50, D439–D444 (2021).

29. M. Mirdita, et al., ColabFold: making protein folding accessible to all. Nat. Methods 19, 679–682 (2022).

30. S. Khan, H. S. Mian, L. E. Sandercock, N. Y. Chirgadze, E. F. Pai, Crystal Structure of the Passenger Domain of the Escherichia coli Autotransporter EspP. J. Mol. Biol. 413, 985–1000 (2011).

31. I. R. Henderson, J. Czeczulin, C. Eslava, F. Noriega, J. P. Nataro, Characterization of pic, a secreted protease of Shigella flexneri and enteroaggregative Escherichia coli. Infect. Immun. 67, 5587–96 (1999).

32. K. Al-Hasani, et al., The sigA Gene Which Is Borne on the she Pathogenicity Island of Shigella flexneri 2a Encodes an Exported Cytopathic Protease Involved in Intestinal Fluid Accumulation. Infect. Immun. 68, 2457–2463 (2000).

33. P. J. Sansonetti, J. Mounier, Metabolic events mediating early killing of host cells infected by Shigella flexneri. Microb. Pathog. 3, 53–61 (1987).

34. K. A. Datsenko, B. L. Wanner, One-step inactivation of chromosomal genes in Escherichia coli K-12 using PCR products. Proc. Natl. Acad. Sci. 97, 6640–6645 (2000).

35. M. Meza-Segura, et al., SepA Enhances Shigella Invasion of Epithelial Cells by Degrading Alpha-1 Antitrypsin and Producing a Neutrophil Chemoattractant. Mbio 12, e02833–21 (2021).

36. Z. Benjelloun-Touimi, P. J. Sansonetti, C. Parsot, SepA, the major extracellular protein of Shigella flexneri: autonomous secretion and involvement in tissue invasion. Mol. Microbiol. 17, 123–135 (1995).

37. I. R. Henderson, J. Czeczulin, C. Eslava, F. Noriega, J. P. Nataro, Characterization of Pic, a Secreted Protease of Shigella flexneri and Enteroaggregative Escherichia coli. Infect. Immun. 67, 5587–5596 (1999).

38. S. A. Bustin, et al., The MIQE Guidelines: Minimum Information for Publication of Quantitative Real-Time PCR Experiments. Clin. Chem. 55, 611–622 (2009).

39. D. Bouyssié, et al., Proline: an efficient and user-friendly software suite for large-scale proteomics. Bioinformatics 36, 3148–3155 (2020).

