## Supplementary figures for "SPATEs promote the survival of *Shigella* to the plasma complement system upon local hemorrhage and bacteremia"

**This PDF file includes:**

Figures S1 to S6
Tables S1 to S2
SI References

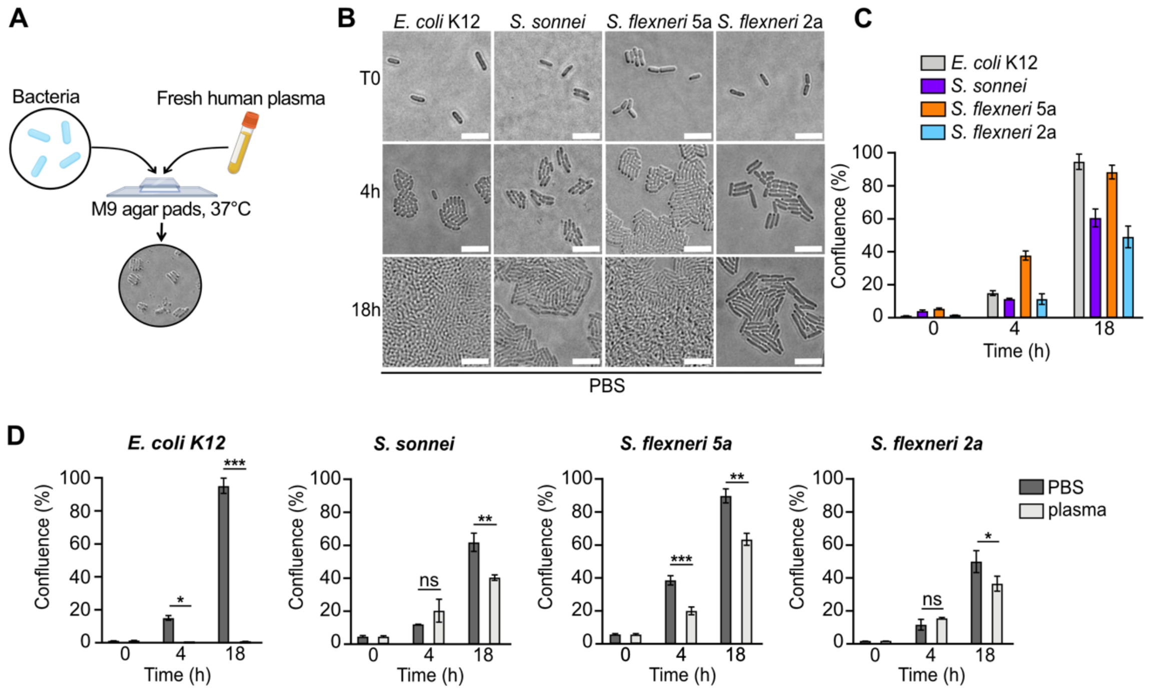


**Figure S1. Comparative analysis of bacterial growth in PBS and plasma.**
**(A)** Experimental procedure used to study bacterial growth and survival on M9 agar pads **(B)** *E. coli* K12 and indicated *Shigella* strains were grown on M9 agar pads supplemented with PBS for up to 18h at 37°C. Bars, 2μm. **(C)** Quantification of bacterial growth (B) by calculating confluence (%) of cultures. Results are expressed as mean ± S.D (n=3). **(D)** Comparative analysis of bacterial growth on M9 agar pads upon supplementation of PBS or fresh human plasma. Results are expressed as mean ± S.D (n=3). 'ns' indicates t-test $p > .05$, * indicates t-test $p < .05$, ** indicates t-test $p < .01$, *** indicates t-test $p < .001$.

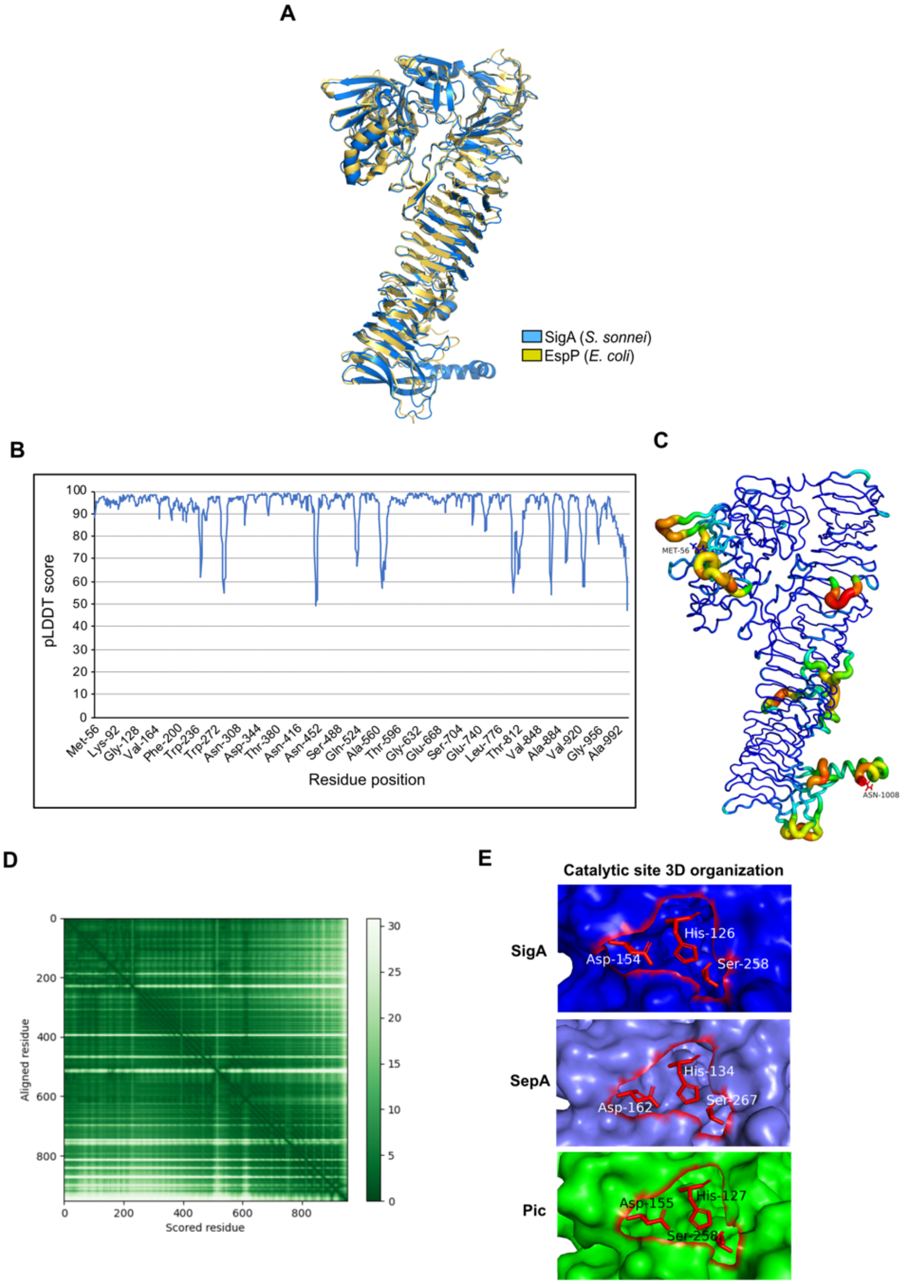


**Figure S2. SigA 3D model prediction and catalytic sites**

**(A)** SigA and EspP passenger domain 3D structure superposition. SigA and EspP (PDB: 3SZE) are colored in blue and gold, respectively. **(B)** SigA AlphaFold 2 pLDDT scores along the sequence. The larger the score the higher the prediction quality. Above 90: very high quality, between 90 and 70: high quality, between 70 and 50: low quality, below 50: very low quality. **(C)** pLDDT scores mapped onto the SigA structure model. A color gradient and thickness of cartoon shapes are associated with structure prediction reliability. Dark blue and thin cartoon regions show high confidence regions while dark red and thick cartoons display less reliable model. **(D)** SigA 3D model predicted aligned error (PAE). The graph shows expected inter-residue distance error, measured in angstroms. On the X-axis, the protein sequence is represented from the N-terminus to the C-terminus from left to right while on the Y-axis it is represented from top to bottom. **(E)** 3D representation of SigA, SepA, Pic catalytic sites. Catalytic residues were drawn as red sticks.

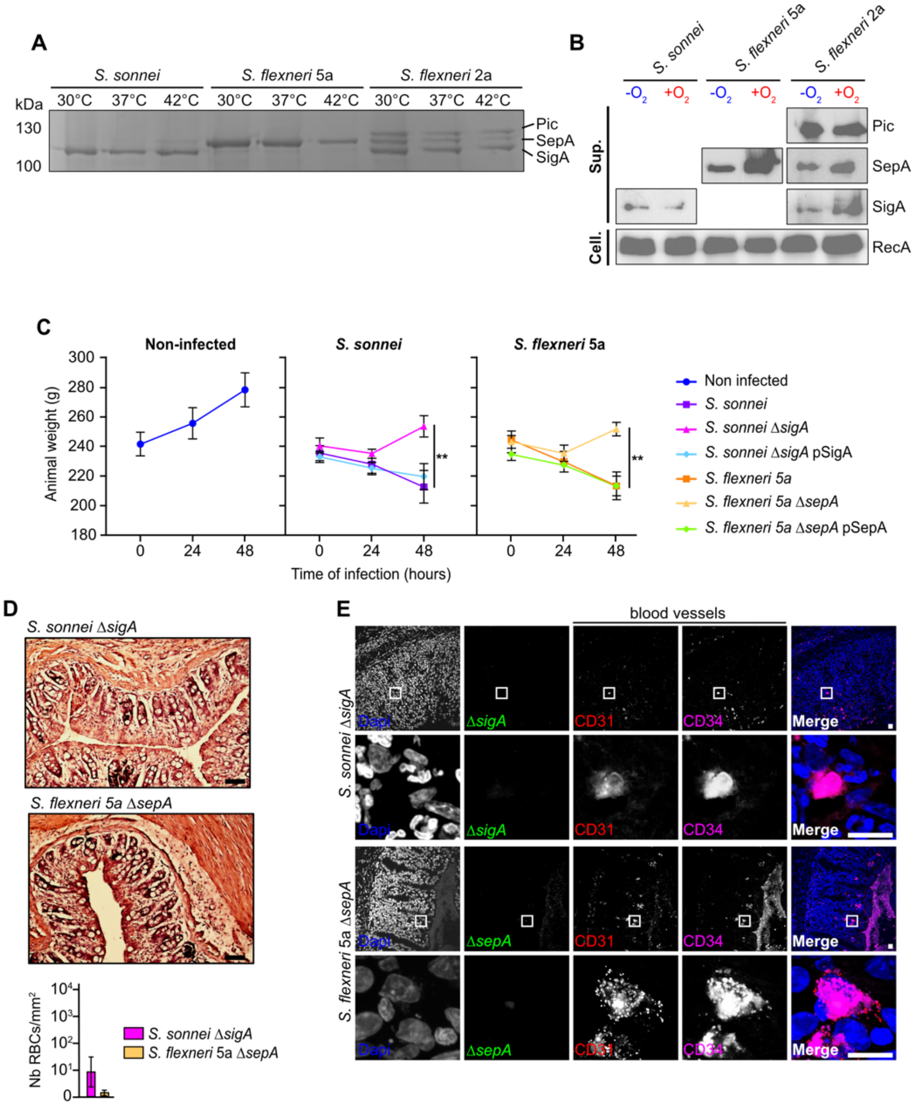


**Figure S3. Regulation of SPATEs secretion and importance during *Shigella* infection**

**(A)** SPATE-containing culture supernatants from *S. sonnei*, *S. flexneri* 5a and *S. flexneri* 2a grown for 5h at 30°C, 37°C and 42°C were separated on 10% SDS-PAGE gel and stained with Coomassie. **(B)** Confirmatory results to Fig. 2C. Western Blot analysis of SPATE-containing culture supernatants (Sup.) from *S. sonnei*, *S. flexneri* 5a and *S. flexneri* 2a grown in -$O_2$ and +$O_2$ conditions. RecA was used as a control (cell.). Representative result of 3 independent experiments. **(C)** Complementary experiment to Fig. 2E. Weight of guinea pigs infected with indicated *Shigella* strains (wild-type, SPATE mutants and complemented strains) for 48 hours. Results are expressed as mean ± S.D (n=3). ** indicates t-test $p$ < .01. **(D)** Complementary result to Fig. 1A. Guinea pig colonic mucosa infected for 48h with *S. sonnei* Δ*sigA* and *S. flexneri* 5a Δ*sepA* strains. Bars, 100 µm. Red Blood Cells (RBCs) which were counted (Nb RBCs/mm$^2$) in each condition. Results are expressed as mean ± S.D (n=3). **(E)** CD31+/CD34+ colonic blood vessels were immunodetected in each condition (red/magenta), together with *S. sonnei* Δ*sigA* and *S. flexneri* 5a Δ*sepA mutant strains* (green). DNA was stained with Dapi (blue). White boxes indicate individual blood vessels. Bars, 20 µm.

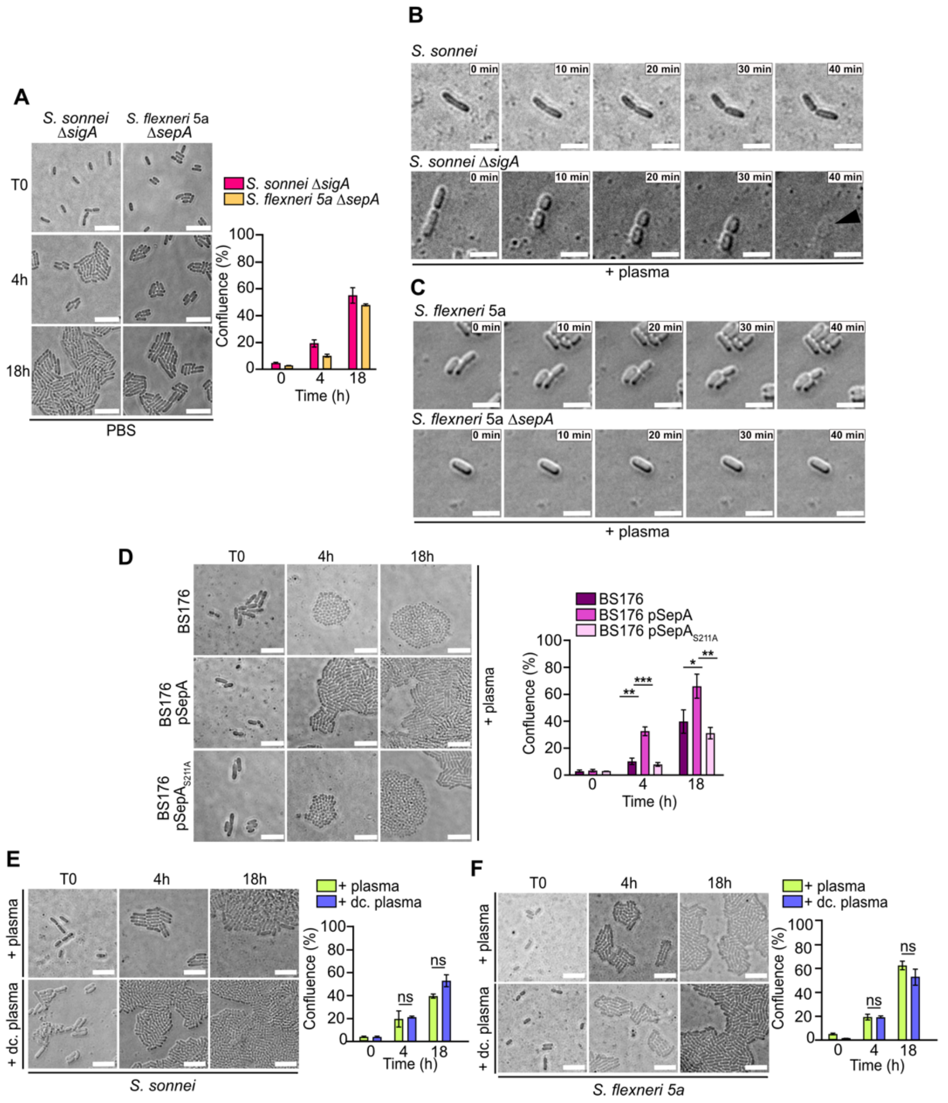


**Figure S4. Importance of SPATE on *Shigella* survival to plasma exposure**

**(A)** *S. sonnei* ΔsigA and *S. flexneri* 5a ΔsepA mutant strains were grown up to 18 h in the presence of PBS on M9 agar pads at 37°C. Quantification of bacterial growth (A) by calculating confluence (%) of cultures. Results are expressed as mean ± S.D (n=3). **(B-C)** Time-course analysis of the growth and survival of (B) *S. sonnei* wild-type and Δ*sigA* mutant strains, and (C) *S. flexneri* 5a wild-type and Δ*sepA* mutant strains on M9 agar pads in the presence of plasma at 37°C for 40 min. Black arrow indicates bacterial lysis. Pictures were acquired every 10 min. Bars, 2 μm. **(D)** Growth of BS176, BS176 pSepA and BS176 pSepA$_{S211A}$ strains on M9 agar pads in the presence of plasma at 37°C for up to 18h. Quantification of bacterial growth (D) by calculating confluence (%) of cultures. Results are expressed as mean ± S.D (n=3). * indicates t-test $p < .05$, ** indicates t-test $p < .01$, *** indicates t-test $p < .001$. **(E-F)** Complementary result to Fig. 3C-D. (E) *S. sonnei* and (F) *S. flexneri* 5a wild-type strains were incubated in the presence of fresh human plasma or decomplemented plasma (dc. plasma) at 37°C for up to 18h. The confluence of cultures was quantified. Results are expressed as mean ± S.D (n=3). 'ns' indicates t-test $p > .05$.

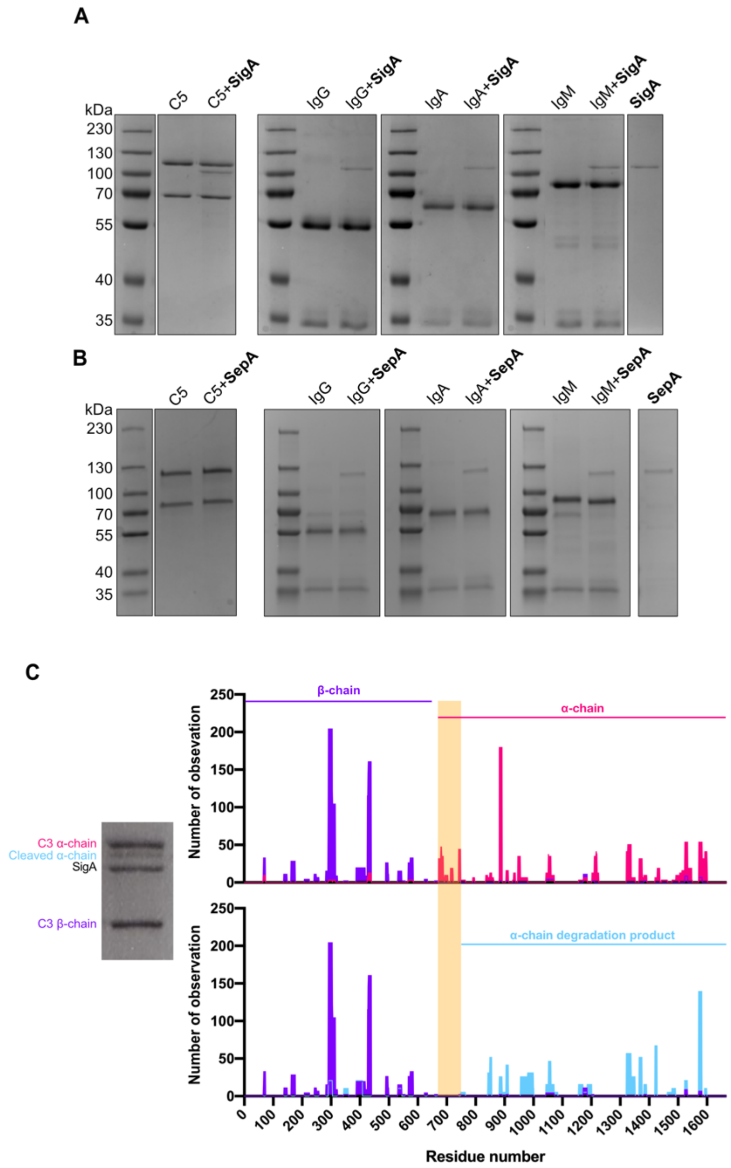


**Figure S5. SigA and SepA substrate specificity and identification of SigA cleavage site on complement component 3 (C3)**

**(A)** Commercially purified human complement component 5 (C5), immunoglobulin G (IgG), immunoglobulin A (IgA) and immunoglobulin M (IgM) were incubated with purified SigA or SepA at 37°C overnight. Samples were separated on a 10% SDS-PAGE gel and stained with Coomassie. **(B)** Complementary result to Fig. 3G. Human complement 3 component (C3) was incubated with purified SigA overnight at 37°C and proteins were separated on a 10% SDS-PAGE gel and stained with Coomassie (left panel). Each indicated band (C3 α-chain/β-chain/cleaved α-chain) was analyzed by mass spectrometry. Purple residues correspond to the C3 β-chain, pink residues correspond to the C3 α-chain, and light blue residues correspond to the cleaved C3 α-chain. The orange area corresponds to the site of cleavage of C3 by SigA.

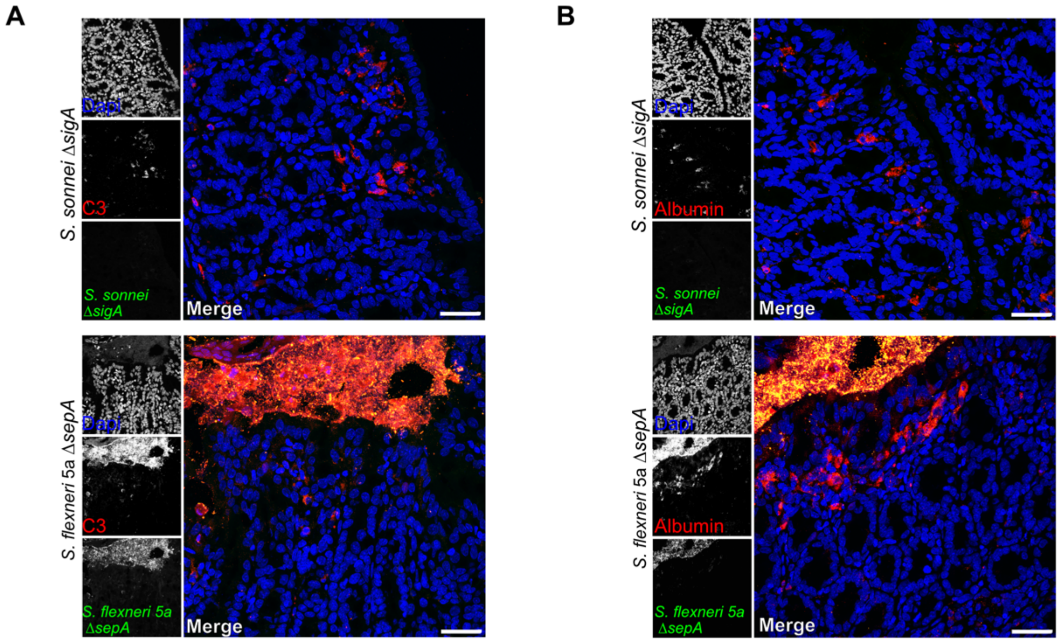


**Figure S6. Complementary result to Fig. 4.**
**(A-B)** Guinea pig colonic mucosa infected by *S. sonnei* Δ*sigA* and *S. flexneri* 5a Δ*sepA* strains (green) for 48h. Infected tissues were stained with an (A) anti-human complement C3 antibody or (B) an anti-human albumin antibody (red) and DNA was stained with Dapi. Bars, 50 μm.

### Table S1 List of strains and plasmids

| Strains | Description | Antibiotic resistance/inductor | Origin |
|---|---|---|---|
| *E. coli* K12 | WT strain | | Lab collection |
| *S. flexneri* 2a | WT strain | | Lab collection |
| *S. flexneri* 5a | WT strain | | Lab collection |
| BS176 | Virulence plasmid-cured *S. flexneri* 5a | | (1) |
| *S. sonnei* CNRS | Clinical isolate CIP106347 | Streptomycin | Pasteur institute collection |
| *S. sonnei* Δ*sigA* | *S. sonnei* CNRS with *sigA* deletion | Streptomycin/Chloramphenicol | Lab collection |
| *S. flexneri* 5a Δ*sepA* | *S. flexneri* 5a with *sepA* deletion | Streptomycin/Chloramphenicol | Lab collection |
| *S. sonnei* Δ*sigA* pSigA3 | *S. sonnei* Δ*sigA* mutant strain complemented with pSigA3 | Streptomycin/Chloramphenicol /Ampicillin IPTG inductor | Lab collection |
| *S. flexneri* 5a Δ*sepA* pSepA | *S. flexneri* 5a Δ*sepA* strain complemented with pZK15 | Chloramphenicol/Ampicillin IPTG inductor | Lab collection |
| BS176 pSepA | *S. flexneri* 5a virulence plasmid cured carrying pZK15 | Ampicillin IPTG inductor | Lab collection |
| BS176 pSepA$_{S211A}$ | *S. flexneri* 5a virulence plasmid cured carrying pSepA$_{S211A}$ | Ampicillin IPTG inductor | Lab collection |
| *E. coli* HB101 pPic1 | *S. flexneri* 5a virulence plasmid cured carrying pPic1 | Tetracyclin | (2) |
| **Plasmid** | **Description** | **Antibiotic resistance/inductor** | **Origin** |
| pKD46 | Plasmid carrying lambda red system | Ampicillin | (3) |
| pKD3 | Plasmid carrying Cm resistance box | Chloramphenicol | (3) |
| pZK15 | Plasmid carrying the *sepA* gene | Ampicillin | (4) |
| pS211A | Plasmid carrying the *sepA* gene with point mutation of 211 catalytic serine in alanine | Ampicillin | Mario Meza-Segura (5) |
| pSigA3 | Plasmid carrying the *sigA* gene | Ampicillin IPTG | Mario Meza-Segura (5) |
| pPic | Plasmid carrying the *pic* gene | Tetracyclin | (2) |
| pGFP | Plasmid carrying the *gfp* gene, with constitutive expression (pFPV25.1) | Ampicillin | Lab collection |

**Table S2. Primers used in this study**

| Primer | Description | Sequence (5’ à 3’) | Size (bp) |
|---|---|---|---|
| ΔsigAsonFor1 | Primers used for the deletion of *S. sonnei sigA* gene | GCTATCCCATAACCACAACTCAGAAATATCGGAGTTCACG T**GTGTAGGCTGGAGCTGCTTC** | 61 |
| ΔsigAsonRev1 | | CTACGGTAGAAGAAGGGCCGCAAACGCGGCCCGGGCTG TTA**CATATGAATATCCTCCTTAG** | 61 |
| ΔsepA5aFor1 | Primers used for the deletion of *S. flexneri* 5a *sepA* gene | CCTATGTAATTAATCTTTGTCAAAATTAGGTTGATGTTTCTA T**GTGTAGGCTGGAGCTGCTTC** | 62 |
| ΔsepA5aRev1 | | CAGAAAAAGGCCTGCCGAATGGCAGGCCTATCCCATTCG **CATATGAATATCCTCCTTAG** | 59 |
| ΔsigAsonk1 | Primers used for the control of *sigA* and *sepA* deletion | GACCGGAAACAACAAAGATC | 20 |
| ΔsigAsonk2 | | GTGATGGCTTCCATGTCGGC | 20 |
| ΔsigAsonk3 | | GGCATGATGAACCTGAATCG | 20 |
| ΔsigAsonk4 | | GCGATATAGTCTGTCACAGG | 20 |
| seQsepAfor1 | | CCAGTCGGCAAAACTAGTTG | 20 |
| seQsepArev1 | | CCAAACTGCCCCTTATCGATACCG | 24 |
| qPCRrrsAF | qRT-PCR *rrsA* | AACGTCAATGAGCAAAGGTATTAA | 24 |
| qPCRrrsAR | | TACGGGAGGCAGCAGTGG | 18 |
| qPCRgyrBF | qRT-PCR *gyrB* | GCAAGCCACGCAGTTTCTC | 19 |
| qPCRgyrBR | | GCTGGTCAGCGAACTGAACG | 20 |
| qPCRsepAF | qRT-PCR *sepA* | GGTTATTCTTACGTCTGTTGCAGC | 24 |
| qPCRsepAR | | CCATCGGGGCTTTATCAAGTTTACC | 25 |
| qPCRsigAF | qRT-PCR *sigA* | GCTGTTTCTGAACTGACCCGG | 21 |
| qPCRsigAR | | GCACCCGGTCTGAACTCTCC | 20 |
| qPCRpicF | qRT-PCR *pic* | CGCCTCAGTATATCGTCAGC | 20 |
| qPCRpicR | | TACCCACCCGATAAAAAGCG | 20 |

**SI References**


1. P. J. Sansonetti, J. Mounier, Metabolic events mediating early killing of host cells infected by Shigella flexneri. *Microb. Pathog.* **3**, 53–61 (1987).

2. F. Ruiz-Perez, *et al.*, Serine protease autotransporters from Shigella flexneri and pathogenic Escherichia coli target a broad range of leukocyte glycoproteins. *Proc. Natl. Acad. Sci.* **108**, 12881–12886 (2011).

3. K. A. Datsenko, B. L. Wanner, One-step inactivation of chromosomal genes in Escherichia coli K-12 using PCR products. *Proc. Natl. Acad. Sci.* **97**, 6640–6645 (2000).

4. Z. Benjelloun-Touimi, P. J. Sansonetti, C. Parsot, SepA, the major extracellular protein of Shigella flexneri: autonomous secretion and involvement in tissue invasion. *Mol. Microbiol.* **17**, 123–135 (1995).

5. M. Meza-Segura, *et al.*, SepA Enhances Shigella Invasion of Epithelial Cells by Degrading Alpha-1 Antitrypsin and Producing a Neutrophil Chemoattractant. *Mbio* **12**, e02833-21 (2021).
